# Analysis of common and rare *VPS13C* variants in late onset Parkinson disease

**DOI:** 10.1101/705533

**Authors:** Uladzislau Rudakou, Jennifer A. Ruskey, Lynne Krohn, Sandra B. Laurent, Dan Spiegelman, Lior Greenbaum, Gilad Yahalom, Alex Desautels, Jacques Y. Montplaisir, Stanley Fahn, Cheryl H. Waters, Oren Levy, Caitlin M. Kehoe, Sushma Narayan, Yves Dauvilliers, Nicolas Dupré, Sharon Hassin-Baer, Roy N. Alcalay, Guy A. Rouleau, Edward A. Fon, Ziv Gan-Or

## Abstract

**Objective:** We aimed to study the role of coding *VPS13C* variants in a large cohort of late-onset PD (LOPD) patients.

**Methods:** *VPS13C* and its untranslated regions were sequenced using targeted next-generation sequencing in 1,567 PD patients and 1,667 controls from 3 cohorts. Association tests of rare potential homozygous and compound heterozygous variants and burden tests for rare heterozygous variants were performed. Common variants were analyzed using logistic regression adjusted for age and sex in each of the cohorts, followed by a meta-analysis.

**Results:** No bi-allelic carriers of rare *VPS13C* variants were found among patients and two carriers of compound heterozygous variants were found in two controls. There was no statistically significant burden of rare (MAF<1%) or very rare (MAF<0.1%) coding *VPS13C* variants in PD. A *VPS13C* haplotype including the p.R153H-p.I398I-p.I1132V-p.Q2376Q variants was nominally associated with a reduced risk for PD (meta-analysis of the tagging SNP p.I1132V (OR=0.48, 95%CI=0.28-0.82, *p*=0.0052). This haplotype was not in linkage disequilibrium (LD) with the known genome-wide association study (GWAS) top hit.

**Conclusions:** Our results do not support a role for rare heterozygous or bi-allelic *VPS13C* variants in LOPD. Additional genetic replication and functional studies are needed to examine the role of the haplotype identified here associated with reduced risk for PD.

## Introduction

The vacuolar protein sorting 13C (*VPS13C*) gene is located within a risk locus for Parkinson Disease (PD), reported in large genome-wide association studies (GWAS) of European population ^1, 2^. The SNP reported in the GWAS (rs2414739, chr15.hg19:g.61994134G>A) was also studied in several Asian populations and in Iranians, with conflicting results ^3–7^, possibly due to ethnicity-related differences.

Subsequently, homozygous and compound-heterozygous *VPS13C* mutations were identified as a rare cause of early onset PD (EOPD) characterized by rapid progression and early cognitive dysfunction. It was demonstrated that *VPS13C* is partially localized at the mitochondrial membrane, and its silencing led to mitochondrial dysfunction and increased PINK1/Parkin-dependent mitophagy ^8^. A follow-up study in 80 EOPD patients identified an additional patient with compound-heterozygous mutations with similar clinical features to the previously reported patients ^9^. In another recent study, a homozygous deletion in *VPS13C* was reported to be the probable cause of an early onset parkinsonism in one patient ^10^. Thus far, full sequencing studies of *VPS13C* have not been reported in late onset PD (LOPD).

To further study the potential role of *VPS13C* variants in PD, we sequenced its coding and regulatory regions using targeted next-generation sequencing in three cohorts of PD (predominantly LOPD) and in controls. We examined the association of common, rare and bi-allelic *VPS13C* variants on the risk for PD. We further tested whether any coding variant or variants in the untranslated regions of *VPS13C* are in linkage disequilibrium (LD) with the top *VPS13C*-associated GWAS hit, to determine if any of these variants can explain the GWAS association of this locus.

## Methods

### Study population

Three cohorts, with a total of 1,567 unrelated PD patients and 1,667 controls, were included in this study, detailed in Table 1. First cohort was composed of French and French-Canadian participants recruited in Quebec (Canada) and in France. This cohort was previously genotyped using the GWAS OmniExpress array, the ethnicity was confirmed using principal component analysis, and samples that were of different ethnicities were not included in this study. The second cohort was recruited in New York - Columbia University, and was previously described^11^. The majority of participants from New York are of European descent and 38% are Ashkenazi Jewish (AJ), 40% of PD patients and 35% of controls. This difference was not statistically significant, yet we adjusted for ethnicity when analyzing this cohort, to avoid effects of ethnicity on the results. The third cohort was recruited in Israel (Sheba Medical Center) and all participants included in this study from the Israeli cohort are of full AJ origin (all four grandparents are full AJ). All patients were consecutively recruited through the clinics, and they represent the typical LOPD patient population with AAO of about 60 (Table 1), as opposed to the studies published so far on *VPS13C* in EOPD. As detailed below, due to the differences in age and sex (Table 1), statistical analysis was adjusted and included age and sex as co-variates. To account for different ethnicities in the New York cohort, an ethnicity covariate was also introduced in this cohort (GWAS data was not available for this cohort, therefore the reported ethnicity was used and not principal components). All three cohorts were sequenced in the same lab (McGill University), following the same protocol. All PD patients were diagnosed by movement disorder specialists according to the UK brain bank criteria ^12^, without excluding patients with family history of PD, since it is now known that there are familial cases of PD, so patients who reported family history of PD were included. However, it is important to emphasize that in the current study only unrelated patients were included, there were no multiple cases from the same family.

**Table 1:** Study population details.

| Sequenced |  | Cohort | Min Depth of Coverage | Analyzed |  |  |  |  |  |  |
| --- | --- | --- | --- | --- | --- | --- | --- | --- | --- | --- |
| N<br>(cases) | N<br>(controls) |  |  | Cases |  |  | Controls |  |  | Genotyping<br>call rate<br>% |
|  |  |  |  | Mean Age |  |  | Mean age |  |  |  |
|  |  |  |  | N | (SD) | %Male | N | (SD) | %Male |  |
|  |  |  |  | y |  |  | y |  |  |  |
| 543 | 866 | French/French- | 15x | 534 | 59.7(11.4) | 63.7 | 858 | 42.0(13.6) | 51.6 | 99.6 |
|  |  | Canadian | 50x | 498 | 59.8(11.2) | 63.5 | 785 | 42.3(13.5) | 52.0 | 99.2 |
| 533 | 270 | New York | 15x | 520 | 59.3(11.8) | 64.0 | 262 | 65.0(9.8) | 34.0 | 99.9 |
|  |  |  | 50x | 502 | 59.3(11.8) | 64.1 | 245 | 65.4(9.6) | 32.7 | 99.5 |
| 491 | 531 | Israel | 15x | 482 | 60.6(11.7) | 61.8 | 488 | 33.9(7.2) | 57.8 | 99.3 |
|  |  |  | 50x | 478 | 60.7(11.7) | 61.7 | 487 | 33.9(7.2) | 57.7 | 99.3 |

N, number; Min, minimum; y, year

### Standard Protocol Approvals, Registrations, and Patient Consents

The institutional review board (McGill University Health Center Research Ethics Board - MUHC REB) approved the study protocols (reference number IRB00010120). Informed consent was obtained from all individual participants before entering the study.

### DNA extraction and *VPS13C* sequencing

DNA was extracted using a standard salting out protocol. The coding sequence and regulatory regions of *VPS13C* were targeted using molecular inversion probes (MIPs), that were designed as previously described ^13^. MIPs were selected based on their predicted coverage, quality and overlap. All MIPs used to sequence *VPS13C* in the present study are included in Table e-1. Targeted DNA capture and amplification was done as previously described ^14^, and the full protocol is available upon request. The library was sequenced using Illumina HiSeq 2500 platform at the McGill University and Genome Quebec Innovation Centre. Reads were mapped to the human reference genome (hg19) with Burrows-Wheeler Aligner ^15^. Genome Analysis Toolkit (GATK, v3.8) was used for post-alignment quality control and variant calling ^16^, and ANNOVAR was used for annotation ^17^. Data on the frequency of each *VPS13C* variant were extracted from the public database Genome Aggregation Database (GnomAD) ^18^. Validation of the tagging variant p.I1132V was performed using Sanger sequencing, with the following primers: forward 5’ – CCGGGAAGGTAATGACAAAA – 3’, reverse 5’ – CCCCTGATTGAAAAGTCACA– 3’

### Quality control

During quality control (QC) filtration using PLINK software v1.9 ^19^, SNPs with genotyping rate lower than 90% were excluded. Genotyping rate cut-off for individuals was 90%, and individuals with a lower genotyping rate were excluded. SNPs that deviated from Hardy-Weinberg equilibrium set at *p*=0.001 threshold were filtered out. Threshold for missingness difference between cases and controls was set at *p*=0.05 and the filtration script adjusted it with Bonferroni correction. After these QC steps, cohort composition was as described in Table 1. To be included in the analysis, minimum quality score (GQ) was set to 30. Rare variants (minor allele frequency, MAF<0.01 or 0.001) had to have a minimal coverage of >50x to be included, and common variants had to have a minimal coverage of >15x to be included in the analysis.

### Statistical Analysis

The association between common *VPS13C* variants and PD was examined by using logistic regression models using PLINK v1.9, with the status (patient or control) as a dependent variable, age and sex as covariates in all cohorts, and AJ ancestry as an additional covariate in the New York cohort. To analyze rare variants (MAF < 0.01) and very rare variants (MAF < 0.001), an optimised sequence Kernel association test (SKAT-O, R package) was performed ^20^. In addition, we examined using SKAT-O the burden of predicted pathogenic variants with Combined Annotation Dependent Depletion (CADD) score of ≥ 12.37 representing the top 2% of potentially deleterious variants. The effects of SNP genotypes on the AAO was tested using analysis of variance (ANOVA; in R software). Meta-analysis of common variants in the three cohorts was performed using Metafor Package in R software ^21^. Linkage disequilibrium in our data was examined by PLINK v1.9 and LD between discovered SNPs and the GWAS top hit rs2414739 was tested using LDlink application, selecting all non-Finish Europeans ^22^.

### Data availability statement

Anonymized data is available upon request by any qualified investigator.

## Results

### Rare *VPS13C* variants and homozygous or compound heterozygous *VPS13C* variants are not associated with late onset PD

The average coverage of *VPS13C* with the MIPs was 94% of nucleotides covered at > 10x, and 90% covered at > 50x. This coverage, while not ideal, is better than the whole exome sequencing coverage reported in the original paper on *VPS13C* in PD, and better than the whole exome and whole genome sequencing coverage of this specific gene in gnomAD (https://gnomad.broadinstitute.org/). There were no differences in coverage between the cohorts and between patients and controls. A total of 60 rare variants that are either nonsynonymous, stop variants or potentially affect a splicing site were identified in the three cohorts and are detailed in Table e-2.

In order to examine whether rare homozygous or compound heterozygous *VPS13C* variants may cause LOPD, and since patients with *VPS13C* bi-allelic mutations are very rare, we included in this analysis only rare variants with allele frequency < 0.001. Only two carriers of two heterozygous variants were identified, and both were controls (Table e-3), suggesting that bi-allelic mutations are not common in LOPD. Of note, we did not examine whether these two variants were on the same allele or different allele (compound heterozygous), since they were found only in controls, which suggests that rare bi-allelic variants are not involved in LOPD in our cohorts.

To further study a potential role for rare (allele frequency < 0.01) or very rare (allele frequency < 0.001) *VPS13C* nonsynonymous or splice variants in LOPD, a SKAT-O was performed on the variants detailed in Table e-2. In the French and French Canadian cohort, 33 (6.6%) PD patients carried a rare variant compared to 49 (6.2%) in controls. In the NY cohort, 43 PD patients (8.6%) carried a rare variant compared to 29 (11.8%) controls. In the Israeli cohort 57 (11.9%) PD patients carried a rare variant compared to 59 (12.1%) among controls. There was no association between rare variants (French/French Canadian cohort *p*=0.44, New York cohort *p*=0.34, Israel cohort *p*=0.91) or very rare variants (French/French Canadian cohort *p*=0.17, New York cohort *p*=0.85, Israel cohort *p*=0.89) and PD. We further examined whether rare variants that are predicted to be deleterious based on CADD score ≥ 12.37 are enriched in PD (the variants included in this analysis are detailed in Table e-4), and no association was found (French/French Canadian cohort *p*=0.58, New York cohort *p*=0.39, Israel cohort *p*=0.40).

### A *VPS13C* haplotype including the p.R153H-p.I398I-p.I1132V-p.Q2376Q coding variants is nominally associated with reduced risk for PD

We have identified 14 common coding variants in our cohort of French and French Canadians, and 13 such variants in each of NY and Israeli cohorts. More details on the number of carriers and frequencies can be found in Table e-5. To test whether common coding variants in *VPS13C* are associated with LOPD, logistic regression models adjusted for age and sex were performed, and additional adjustment for ethnicity was included in the New York cohort (see methods). A nominal association was observed in four variants (p.R153H [rs12595158, chr15.hg19:g.62316035C>T], p.I398I [rs9635356, chr15.hg19:g.62299603T>G], p.I1132V [rs3784635, chr15.hg19: g.62254989T>C] and p.Q2376Q [rs17238189, chr15.hg19: g.62212781T>C]) with reduced risk for PD in the New York cohort (Table 2). These remained nominally significant with and without including adjustment for ethnicity, suggesting that ethnicity has no role in this association, and only one of them, p. I1132V (OR 0.28, 95% CI 0.12-0.64, *p*=0.0025), remained statistically significant after correction for multiple comparisons. In the two other cohorts, these variants showed the same directionality as in the New York cohort, but did not reach statistical significance (Table 2). These four variants, p.R153H, p.I398I, p.I1132V and p.Q2376Q, are in strong or even complete LD (Table e-6). The most significant tagging SNP of this haplotype, p. I1132V, was validated using Sanger sequencing in all three cohorts. Meta-analysis of all the common coding variants showed an association of these four linked SNPs with reduced risk for PD (Figure 1), with similar directionality across all cohorts. To examine whether this haplotype may affect AAO of PD, ANOVA with the status of the p. I1132V variant was performed. This variant was not associated with AAO in all three cohorts (French/French Canadian cohort *p*=0.65, New York cohort *p*=0.34, Israel cohort *p*=0.99).

**Table 2:**
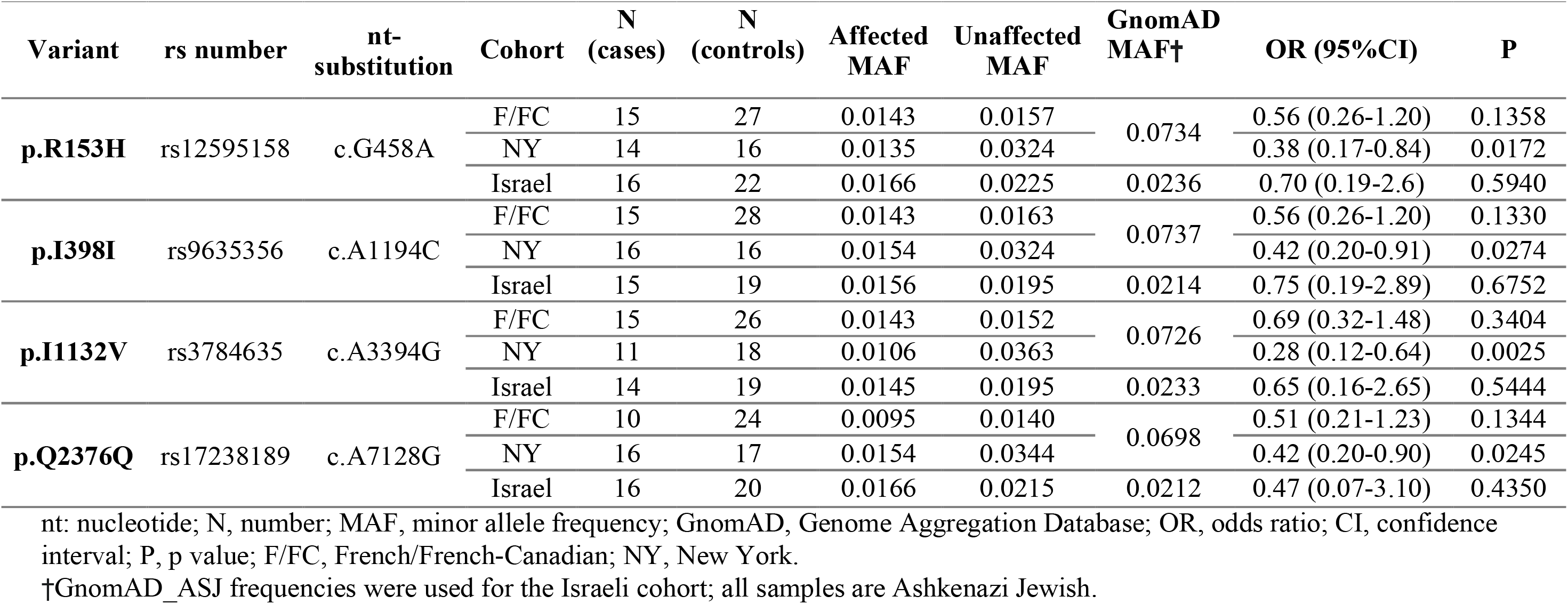
Four variants forming the protective haplotype found in *VPS13C* and the results of the logistic regression in three cohorts.

**Figure 1 Legend:**
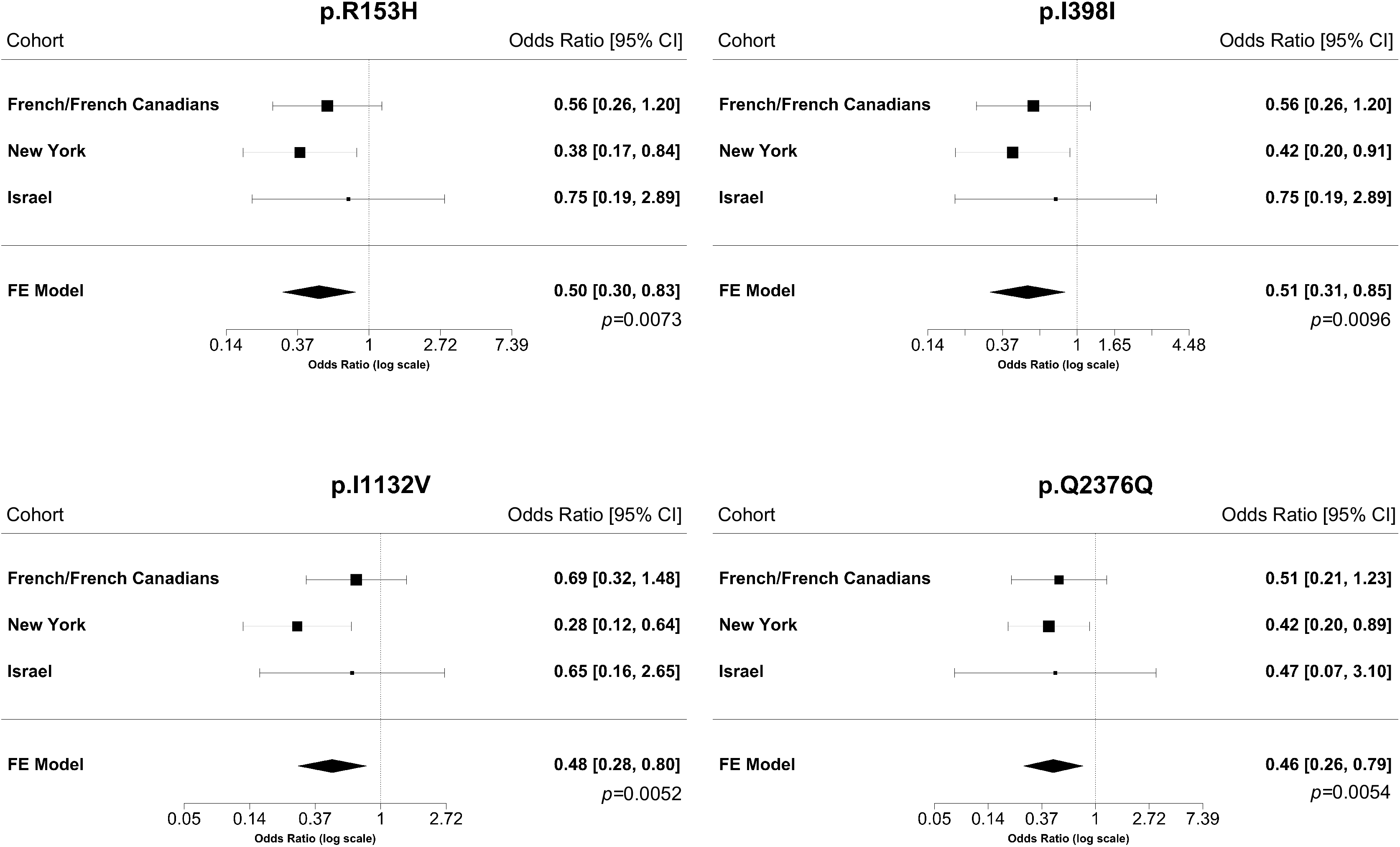
Forest plots – meta-analyses of four *VPS13C* coding variants associated with reduced risk for Parkinson Disease. The forest plots depict the effects of the four *VPS13C* coding variants that create the haplotype associated with reduced PD risk in the three cohorts studied, and their meta-analyzed effect on risk for PD. The results in the random effect model are nearly identical. CI, confidence interval; FE, fixed effect

None of these variants were in LD with the known GWAS top hit, rs2414739, and therefore these associations do not explain the GWAS hit in the *VPS13C* locus. Of all the other common coding variants, only one variant, p.E2008D (rs78071599, chr15.hg19: g.62223303C>G), was in some LD with the GWAS top hit (D’=0.808, r^2^=0.006, Table e-7), but this variant was not associated with LOPD, and therefore it also cannot explain the GWAS hit in our populations. Interestingly, an intronic variant, rs78530361 (chr15.hg19: g.62214265C>T), was in strong LD with the top GWAS SNP (D’=1, r^2^=0.003). Of note, r^2^ is low since this intronic SNP has a much lower allele frequency than the top GWAS hit, but every time the rs78530361 SNP is found, it is on an allele which harbors the top GWAS hit rs2414739. However, in our cohorts, this variant was not associated with PD (*p*=0.66), likely due to its very low r^2^ with the GWAS hit.

## Discussion

Our study, which included full sequencing of *VPS13C* in three cohorts, identified 60 rare *VPS13C* variants (MAF<0.01) that are nonsynonymous or affect splicing, and 18 common variants (MAF>0.01) in coding regions of the gene and splice sites. Our results suggest that rare homozygous and compound heterozygous variants are rare in LOPD and probably have importance mainly in early onset PD, as previously described. We have identified a potentially protective haplotype, which includes four variants, two of which are substitutions of amino acids, p.R153H and p.I1132V. The association was driven by the NY cohort mainly, but the two other cohorts demonstrated similar directionality of effect and effect size, and also contributed to the association. It is unlikely that differences in ethnicity in the NY cohort drove the association, since analysis was with adjustment for ethnicity, and analysis without adjustment yielded nearly identical results, ruling out effect of ethnicity. The other two cohorts were of homogeneous ethnicities, therefore in these cohorts too, ethnicity could not have affected the results. Interestingly, this haplotype is not in LD with the top GWAS SNP, rs2414739, suggesting that this may be a secondary association in the *VPS13C* locus, which was not identified in previous studies. However, this should be considered as preliminary results and needs to be examined in additional cohorts in order to conclude whether this haplotype is indeed associated with reduced risk for PD. Since the disease-causing mutations reported in *VPS13C* are loss-of-function mutations, a protective effect could occur for example due to gain of function or overexpression. In GTEx (https://www.gtexportal.org/home/) these variants were not associated with increased expression or affect splicing. The two nonsynonymous variants of this haplotype, p.R153H and p.I1132V, have high CADD scores (22.8 and 13.7, respectively), suggesting that they may affect the protein structure or function. Whether there is such effect and whether it is associated with gain-of-function will need to be examined in follow-up studies. Furthermore, the full sequencing analysis did not identify any coding variant that is in LD with the original GWAS hit that can explain the GWAS association in this locus. This may suggest that the variant that has the main effect on the risk for PD in the *VPS13C* is outside of the coding and untranslated regions of the gene, likely being a regulatory element.

Previous studies on the top GWAS hit in the *VPS13C* locus demonstrated conflicting results. While significant associations of the top GWAS hit in this locus (rs2414739) with PD were found in Iranian ^5^ and East Asian ^3^ populations, negative results were reported in Taiwanese ^4^ and Han Chinese ^6, 7^ populations. Therefore, it is possible that the association of *VPS13C* with PD is population dependent. Of note, that the association in the current study was mainly driven by populations enriched in Ashkenazi Jews, while in the French/French Canadian cohort the differences between patients and controls were much smaller and not statistically significant (Table 2). This may suggest that this haplotype association is population specific, and additional studies in different populations are required to answer this question.

In our three cohorts, we did not find any very rare homozygous variants with MAF<0.001 and found only two carriers of two heterozygous variants, both of which were controls (Table e-3). We were unable to determine if the variants were in cis or trans, but since no carriers of two variants was found among patients, it is clear that *VPS13C* bi-allelic mutations do not contribute to PD in our cohorts. Previously reported cases of PD in carriers of compound heterozygous or homozygous mutations all shared a specific clinical presentation of PD: early onset, rapid progression and early cognitive dysfunction ^8, 9^. In one study, the patient’s AAO was 39, and disease progression was moderately severe with psychiatric symptoms and impaired cognition ^9^. In another study, the three patients showed severe phenotypes: AAO of 25, 33, and 46 years, severe and early cognitive dysfunction, and became bedridden at 31, 43, and 58 years, respectively. Considering the young age of one of the compound heterozygous *VPS13C* variant carriers in our cohort (30 years), it is still possible, although unlikely that this individual will develop PD in the future. The negative results of the SKAT-O analyses demonstrate that rare heterozygous variants in *VPS13C* do not have an important role in PD in our cohorts.

Our study has several limitations. The differences between PD patients and controls in sex and mainly age are significant in some of our cohorts. To address this limitation, we included age and sex as covariates in the regression models. Therefore, if the association of the protective haplotype was related to age and not to disease status, we would likely not observe an association in the adjusted model. Furthermore, the association with the haplotype had the same directionality and similar effect size in the NY cohort in which the controls were older (and the association was statistically significant), and in the other two cohorts where the controls were younger, likely ruling out effect of age. There was no significant difference in the percentage of Ashkenazi Jews between patients and controls in the New York cohort, yet we still performed the regression model with and without ethnicity as a covariate, and in both analyses the results remained significant. Nevertheless, the fact that our populations are enriched with relatively homogeneous populations such as Ashkenazi Jews and French Canadians requires additional studies in other populations. Another potential limitation is that we could not analyze the effect of CNVs in *VPS13C* with our data. Loss of function and exonic deletions/duplications are rare in gnomAD, found in only 3 individuals at a heterozygous state, and therefore not likely to have a major contribution in PD. However, future studies to examine the potential role of *VPS13C* in PD are required. Furthermore, as no functional experiments were performed in the current study, the potential effects of variants that we report here should be examined in additional studies.

In conclusion, our results suggest that *VPS13C* variants have a limited role in late-onset PD. The potentially PD protective haplotype located within *VPS13C,* which requires additional replications, may suggest that *VPS13C* could be a future target for PD therapeutic development. If naturally occurring genetic variants may reduce PD risk, it is conceivable that drugs that can mimic their effects could be developed. Additional genetic and functional studies will be required to determine if *VPS13C* may be a viable target for PD drug development.

## Acknowledgments

We thank the patients and control subjects for their participation in this study. This work was financially supported by the Michael J. Fox Foundation, the Canadian Consortium on Neurodegeneration in Aging (CCNA) and the Canadian Glycomics Network (GlycoNet). This research was also undertaken thanks in part to funding from the Canada First Research Excellence Fund (CFREF), awarded to McGill University for the Healthy Brains for Healthy Lives (HBHL) program. The Columbia University cohort is supported by the Parkinson’s Foundation, the National Institutes of Health [K02NS080915, and UL1 TR000040] and the Brookdale Foundation. GAR holds a Canada Research Chair in Genetics of the Nervous System and the Wilder Penfield Chair in Neurosciences. EAF is supported by a Foundation Grant from the Canadian Institutes of Health Research (FDN grant – 154301). ZGO is supported by the Fonds de recherche du Québec - Santé (FRQS) Chercheurs-boursiers award, and by the Young Investigator Award by Parkinson Canada. The access to part of the participants for this research has been made possible thanks to the Quebec Parkinson’s Network (http://rpq-qpn.ca/en/). We thank Armaghan Alam, Daniel Rochefort, Helene Catoire, Clotilde Degroot and Vessela Zaharieva for their assistance.

## Appendix 1: Authors

| Name | Location | Role | Contribution |
| --- | --- | --- | --- |
| Uladzislau Rudakou, BSc | McGill University,<br>Montreal, QC, Canada | Author | Analyzed and interpreted the data; performed validations; drafted the manuscript for intellectual content |
| Jennifer A. Ruskey, MSc | McGill University,<br>Montreal, QC, Canada | Author | Major role in the acquisition of data; revised the manuscript for intellectual content |
| Lynne Krohn, BSc | McGill University,<br>Montreal, QC, Canada | Author | Interpreted the data; revised the manuscript for intellectual content |
| Sandra B. Laurent, BSc | McGill University,<br>Montreal, QC, Canada | Author | Major role in the acquisition of data; Interpreted the data; revised the manuscript for intellectual content |
| Dan Spiegelman, MSc | McGill University,<br>Montreal, QC, Canada | Author | Major role in the acquisition of data; revised the manuscript for intellectual content |
| Lior Greenbaum, MD, PhD | Sheba Medical Center,<br>Tel Hashomer, Ramat Gan, Israel | Author | Major role in the acquisition of data; revised the manuscript for intellectual content |
| Gilad Yahalom, MD | Sheba Medical Center,<br>Tel Hashomer, Ramat Gan, Israel | Author | Major role in the acquisition of data; revised the manuscript for intellectual content |
| Alex Desautels, MD, PhD | Université de Montréal,<br>Montreal, QC, Canada | Author | Major role in the acquisition of data; revised the manuscript for intellectual content |
| Jacques Y. Montplaisir, MD, PhD | Université de Montréal,<br>Montreal, QC, Canada | Author | Major role in the acquisition of data; revised the manuscript for intellectual content |
| Stanley Fahn, MD | Columbia University,<br>New York, NY, USA | Author | Major role in the acquisition of data; revised the manuscript for intellectual content |
| Cheryl H. Waters, MD | Columbia University,<br>New York, NY, USA | Author | Major role in the acquisition of data; revised the manuscript for intellectual content |
| Oren Levy,<br>MD, PhD | Columbia University,<br>New York, NY, USA | Author | Major role in the acquisition of data; revised the manuscript for intellectual content |
| Caitlin M.<br>Kehoe, MA | Columbia University,<br>New York, NY, USA | Author | Major role in the acquisition of data; revised the manuscript for intellectual content |
| Sushma<br>Narayan, MS | Columbia University,<br>New York, NY, USA | Author | Major role in the acquisition of data; revised the manuscript for intellectual content |
| Yves<br>Dauvilliers,<br>MD, PhD | University of<br>Montpellier,<br>Montpellier, France | Author | Major role in the acquisition of data; revised the manuscript for intellectual content |
| Nicolas<br>Dupré, MD,<br>FRCPC | Université Laval,<br>Québec City, QC,<br>Canada | Author | Major role in the acquisition of data; revised the manuscript for intellectual content |
| Sharon<br>Hassin-Baer,<br>MD | Sheba Medical Center,<br>Tel Hashomer, Ramat<br>Gan, Israel | Author | Major role in the acquisition of data; revised the manuscript for intellectual content |
| Roy N.<br>Alcalay, MD,<br>MS | Columbia University,<br>New York, NY, USA | Author | Major role in the acquisition of data; revised the manuscript for intellectual content |
| Guy A.<br>Rouleau, MD,<br>PhD, FRCPC,<br>FRSC | McGill University,<br>Montreal, QC, Canada | Author | Major role in the acquisition of data; revised the manuscript for intellectual content |
| Edward A.<br>Fon, MD | McGill University,<br>Montreal, QC, Canada | Author | Major role in the acquisition of data; revised the manuscript for intellectual content |
| Ziv Gan-Or,<br>MD, PhD | McGill University,<br>Montreal, QC, Canada | Author | Designed and conceptualized study; major role in the acquisition of data; assisted with interpretation of the data revised the manuscript for intellectual content; led and coordinated communication among sites. |

